# Distinct functional classes of excitatory neurons in mouse V1 are differentially modulated by learning and task engagement

**DOI:** 10.1101/533463

**Authors:** Joseph B. Wekselblatt, Cristopher M. Niell

**Affiliations:** Institute of Neuroscience and Department of Biology, University of Oregon, Eugene OR 97403

## Abstract

Learning can cause significant changes in neural responses to relevant stimuli, in addition to modulation due to task engagement. However, it is not known how different functional types of excitatory neurons contribute to these changes. To address this gap, we performed two-photon calcium imaging of excitatory neurons in layer 2/3 of mouse primary visual cortex before and after learning of a visual discrimination. We found that excitatory neurons show striking diversity in the temporal dynamics of their response to visual stimuli during the behavior, and based on this we classified them into transient, sustained, and suppressed groups. Notably, these functionally defined cell classes exhibit different visual stimulus selectivity and modulation by locomotion, and were differentially affected by training condition. In particular, we observed a decrease in the number of transient neurons responsive during behavior after learning, while both transient and sustained cells showed an increase in modulation due to task engagement after learning. The identification of functional diversity within the excitatory population, with distinct changes during learning and task engagement, provides insight into the cortical pathways that allow context-dependent neural representations.

## Introduction

Sensory perception is context dependent, and has been shown to be modulated by a number of factors including task demands, learning, and behavioral engagement. Specific task demands can modulate neural responses to improve the detection or discriminability of relevant stimulus features and spatial locations (Deubel et al., 1998; Slotnick et al., 2003; Poort et al., 2015). Likewise, learning can affect coding strategies leading to increasingly sparse or robust representations, changes in tuning, and shifts in excitatory/inhibitory balance (Yizhar et al., 2011; Peters et al., 2014; Chu et al., 2016). Task engagement and arousal have also been associated with a multiplicative gain in response to relevant sensory stimuli reminiscent of studies showing that attention can enhance responses (Moran and Desimone, 1985.; McAdams and Maunsell, 1999; Niell and Stryker, 2010; McGinley et al., 2015; Vinck et al., 2015; Wekselblatt and Niell, 2015).

Circuit mechanisms for context dependent processing of sensory information are poorly understood, especially among subpopulations of excitatory neurons. Recent studies have addressed the differences in the responses of inhibitory neuron subtypes (representing only 10-20% of cortical neurons) and their effects on learning (Kato et al., 2015; Kuchibhotla et al., 2016; Yavorska and Wehr, 2016), finding differential modulation of responses due to training in three genetically distinct subtypes. It remains unclear, however, how much diversity exists in the excitatory population, which represents approximately 80% of the neurons in cortex, and how potential sub-populations of the excitatory pool might contribute to context dependent perception. Furthermore, inhibitory neurons primarily affect local processing, while excitatory neurons, especially those in layer 2/3, send long range projections to other brain areas, determining what information is transmitted downstream (Han et al., 2018). Changes in the response of excitatory cells could greatly impact the types of information relayed to other cortical regions.

In the current study, we sought to test whether excitatory neurons in primary visual cortex showed similar learning related changes across the whole population, or whether changes due to learning were specific to different sub-populations. Furthermore, if the changes are specific, what features of the neural response can define these sub-populations? To determine the changes in excitatory neurons due to learning and context, we employed two-photon imaging in head-fixed animals while they performed a visual discrimination task both before and after learning, or simply after exposure to the stimuli without learning. We clustered the neural responses based on temporal dynamics, rather than visual tuning properties, revealing three distinct cell classes (transient, sustained, suppressed) among the responsive neurons. We refer to these as ‘functional’ cell types, as they are defined by their activity patterns rather than other features such as gene expression or morphology. This diversity in the excitatory population of layer 2/3 also corresponded to specific changes in responsiveness over the course of learning, and as well as modulation by task engagement that was cell type-specific and dependent on learning history. These results reveal striking heterogeneity in the excitatory cell population in cortex, and suggest that changes over learning are more specific than previously described.

## Materials and Methods

### Animal use

Animals were maintained in the animal facility at University of Oregon and used in accordance with protocols approved by the University of Oregon Institutional Animal Care and Use Committee (IACUC). All procedures were conducted in accordance with the ethical guidelines of the National Institutes of Health. Animals were maintained on a 12hr light / 12hr dark reverse light cycle, with training and experiments performed during the dark phase of the cycle. Running wheels were provided in each home cage as environmental enrichment.

### Surgical Procedures

The headplate and cranial window implant procedures were performed as described in Wekselblatt et al. (2016), with minor modifications described below. Briefly, a titanium headplate was cemented to the skull to allow head fixation in an initial surgery. Following recovery from the headplate surgery, a 5mm diameter cranial window (No. 1 coverglass, Warner Instruments) was implanted over the visual cortex (3mm lateral, 1.5mm anterior of Bregma). For the headplate procedure, the imaging well of the headplate was filled with a thin layer of clear dental acrylic to protect the skull, instead of sylgard used in the previous description. We found that dental acrylic provided greater protection, as the sylgard plug would occasionally fall out or cause condensation to be trapped inside the headplate well. For the cranial window procedure, dexamethasone sodium phosphate (2 mg/kg subcutaneously) was administered ~24 hr prior to surgery and again ~2hr prior to surgery to reduce brain edema. We found that giving two doses of dexamethasone, instead of the single dose previously described, had a much greater effect on reducing brain swelling caused by the craniotomy. Finally, mice were allowed to recover for at least 1 week after surgeries before beginning habituation and training. For the first 3 days we provided post-surgical animals with wet food in a small dish on the floor of their cage and a high calorie nutrient supplement (Nutri-Cal, Vetoquinol). This prevented weight loss after surgery and helped animals recover quickly.

### Imaging

We used a custom macroscope for widefield imaging, as described in Wekselblatt et al. (2016). This design was based on the tandem lens system used for intrinsic signal imaging (Ratzlaff and Grinvald, 1991; Kalatsky and Stryker, 2003). Briefly, two camera lenses (Nikon 50mm f/1.2 and Nikon 105mm f/1.8) were interfaced with a dichroic filter cube (Thorlabs). Blue light illumination was supplied in epifluorescence configuration through the filter cube housing to excite the fluorescent calcium indicator. Green light was supplied directly through a fiber placed obliquely above the brain to measure intrinsic signal due to hemodynamics. Images were acquired at 10Hz with 4x spatial binning using Camware software (PCO Corporation), with frame acquisition and LED illumination triggered by TTL pulses from the stimulus presentation computer to synchronize with visual stimulus frames.

Two-photon imaging was performed using a resonant scanning two photon microscope optimized for *in-vivo* imaging (Neurolabware, Los Angeles, CA) coupled to a Mai-Tai HP Ti-Sapphire pulsed laser tuned to 920nm, with a 16x/0.8NA objective (Nikon). Scanbox software (Dario Ringach / Neurolabware) in Matlab (Mathworks) was used for data acquisition. Images were acquired at 10 fps and 796×796 pixels over a ~800×800um field of view, using 65-110mW illumination power as measured at the front aperture of the objective. All recordings were targeted to layer 2/3 of V1 (depth 100 – 200 microns).

### Stimulus delivery and behavior control

Visual stimuli were presented on a Viewsonic VA2342 LCD monitor (28 × 50cm), linearized to correct for gamma (mean luminance 35cd/m^2^), oriented tangentially 25 cm from the mouse’s right eye in portrait configuration (except during the shaping task), covering ~60×90° of visual space. Stimuli were generated with custom software using the Psychtoolbox extension for Matlab (Brainard, 1997; Pelli, 1997).

The behavioral control system was as described in Wekselblatt et al. (2016). Movement of the mouse on the spherical treadmill was measured with an optical USB computer mouse positioned laterally on the polystyrene ball, acquired once per stimulus frame (60Hz) in Matlab. Visual stimuli for behavior consisted of 33° diameter circular patches of stationary square wave gratings at random spatial phase with spatial frequency 0.08cpd, oriented either horizontally or vertically. Gratings appeared at one of two locations on the screen, either top or bottom (centered at +/− 20° from center of monitor in elevation).

For retinotopic mapping, we binarized a 1/f noise stimulus described previously (Pnevmatikakis et al., 2016; Wekselblatt et al., 2016) with spatial frequency corner of 0.05cpd and cutoff of 0.12cpd, and temporal frequency cutoff of 5Hz. This noise stimulus was masked to create a 20 degree wide bar that moved across the visual display with a 10 second period in either azimuth or elevation for 30 cycles. This stimulus was binarized to black/white instead of grayscale to increase contrast and generate edges.

Passive viewing stimuli consisted of static grating patches identical to those presented during behavior (33° diameter, 0.08cycle/degree), with an additional location in the center of the screen and 2 additional orientations (45 and 135 degrees). Passive sessions consisted of 20 repetitions per stimulus, presented for 1 second each with an inter-stimulus interval of 1 second.

### Behavioral training

The spherical treadmill and head-fixation were as described in Wekselblatt et al. (2016). Briefly, a 200mm hollow polystyrene ball was placed inside a 250mm polystyrene hemisphere (Graham Sweets Studios, Cardiff, UK) supplied with airflow through a Tygon tube positioned at the bottom pole. This provided a rotating surface on which the mouse was able to freely move. The animal’s head was fixed via a surgically attached headplate that could be screwed into a rigid crossbar above the floating ball. Headplates were manufactured from titanium by emachineshop.com. Designs are available upon request.

Prior to beginning behavior, mice were handled for several days until they were comfortable with the experimenter. Once water scheduling began, animals received water only during and immediately after head-fixed training on the ball. Training sessions increased in duration over the course of 1–2 weeks, from 30 minutes to 1 hour sessions, and occurred twice per day separated by ~3 hours.

All training was performed with a mostly automated system (Wekselblatt et al., 2016). First, mice learned the simple visual task of discriminating the location of a luminance stimulus with the screen in landscape orientation. Initially, animals were required to request a trial by stopping spontaneous locomotion for 1 second to receive a water reward. Upon requesting a trial, a stimulus was presented with dark on either the nasal or lateral 2/3 of the screen, and light on the other 1/3. The animal was rewarded with a second water drop for moving either right or left on the ball if dark was on the nasal or lateral side respectively. When an animal could reliably request trials, the water reward for stopping to initiate trials was eliminated. This task took about 1-2 weeks to learn and established trial structure and attention to the visual stimulus.

Water rewards were calibrated by the experimenter to maintain consistent weights (>80% of baseline), corresponding to ~0.8 – 1.0 ml of water through the course of a session. Mice were trained 2 sessions per day, 7 days per week for 30 – 60 minutes per session, with a session ending when the subject stopped initiating trials or performance dropped.

Once a mouse could perform a significant number of trials (>200 per session) and reached ~85% accuracy on the luminance task, they were graduated to one of three task conditions: naïve, experienced, or trained (Figure 1C). These tasks followed a similar trial structure to the luminance task, but a circular grating patch, either horizontal or vertical, was presented on either the top or bottom of the screen (Figure 1D). The stimulus remained on the screen for one second after correct responses. Incorrect responses triggered a 3.5sec timeout with potentially aversive flashing error stimulus and 50% probability of a correction trial, where the previous stimulus was repeated until answered correctly, to prevent perseverative errors and bias.

**Figure 1.**
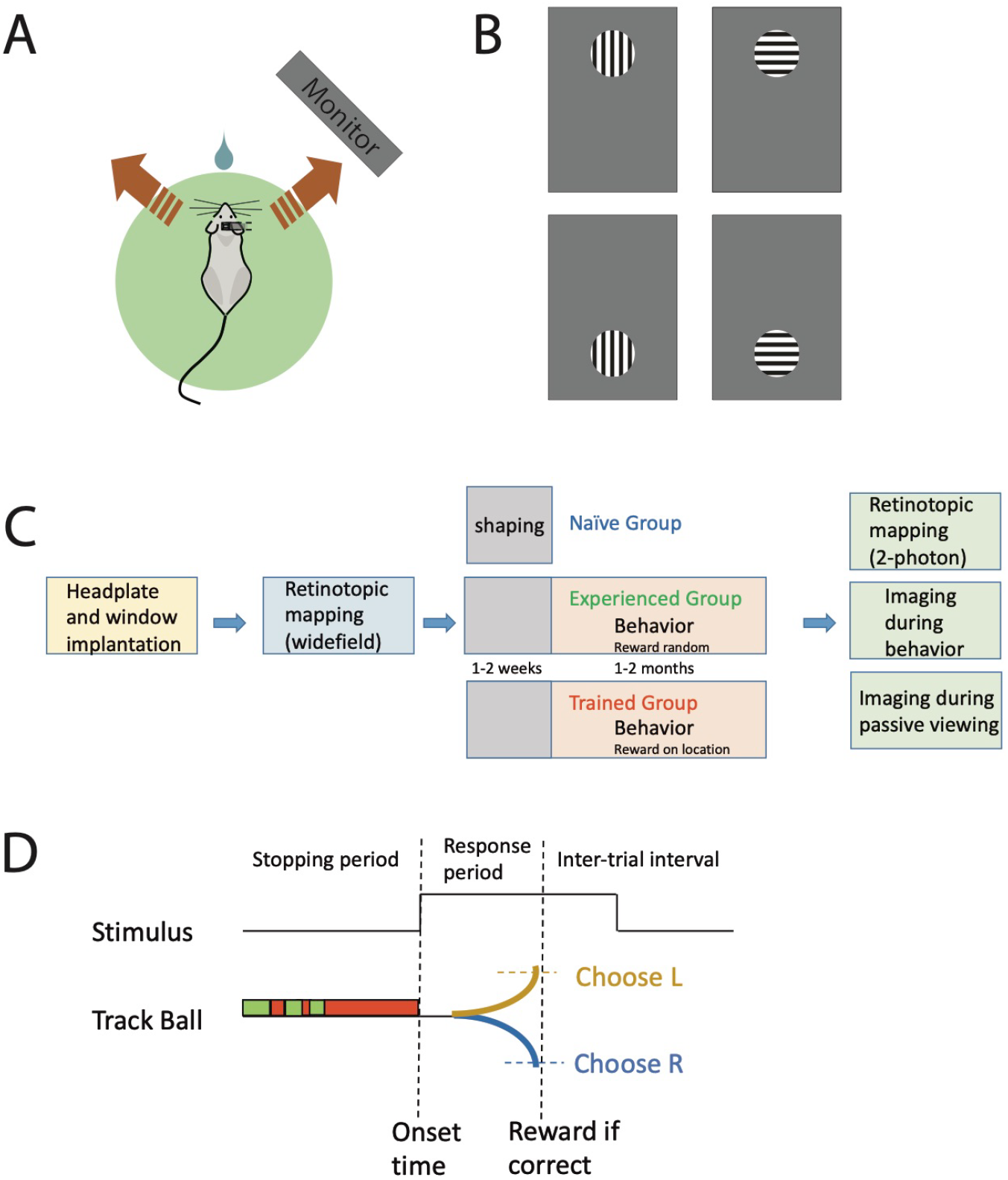
Behavioral paradigm and training. **A)** Schematic of spatial 2-alternative forced choice (2-AFC) visual behaviors. Head-fixed mice move the spherical treadmill a threshold distance right or left in response to the location of stimuli presented on a monitor. **B)** Stimuli presented for visual behaviors were a single square wave grating patch of horizontal or vertical orientation located at the top or bottom of monitor. **C)** Flow chart showing sequence of procedures and behavioral training for each experimental group. **D)** Time course of a behavioral trial. During the stopping period, green indicate locomotion and red indicates the animal has stopped. The animal must stop for 1 full second to trigger stimulus onset.

Animals in the ‘naïve’ group were imaged during their initial exposures (<8) to the grating stimuli immediately following mastery of the shaping task. ‘Naïve’ animals were randomly rewarded during imaging sessions to ensure that they stayed naïve to any rule. Animals who were assigned to the ‘experienced’ group were moved to the grating stimuli but the spatial location of the stimulus did not predict reward. Instead, the rewarded direction of motion of the trackball was random on each trial. Animals in the experienced group were given roughly the same amount of experience with the grating stimuli as the ‘trained’ group before imaging occurred (1-2 months, >50 sessions, >10,000 trials). Animals who were assigned to the ‘trained’ group were moved to the grating stimuli (Figure 1B), and tested based on the location of the grating. Animals became proficient at this task in 2-4 weeks. Imaging occurred only after animals performed >200 trials per session at over 80% correct for at least two consecutive sessions.

The imaging behavioral configuration was identical to training, except the stimulus remained on the screen for 1.5 seconds even after incorrect responses, replacing the flashing error stimulus used during training, to avoid differences in visual input between correct and error trials. Additionally, correction trials were removed during imaging sessions to ensure each trial was independent from the previous answer.

### Optogenetic Silencing

To perform optogenetic silencing we activated GABAergic fast-spiking interneurons (Pv+) expressing channelrhodopsin-2 (ChR2) by illumination with a blue LED (470nm) through a chronic cranial window (Lien and Scanziani, 2013; Burgess et al., 2017). The animals used for optogenetic silencing experiments were trained on the spatial discrimination task until they reached criterion performance (>80% correct, >200trials per session). Experimental animals were double positive for cre-dependent ChR2 and Pv-cre. Control animals were GCaMP expressing animals used in the imaging experiments. We provided light stimulation through a chronic cranial window on 20% of trials, starting at visual stimulus onset and lasting until behavioral response. We performed silencing by illuminating either the entire cranial window (which includes several extrastriate areas) or restricting illumination to primary visual cortex (Poort et al., 2015). V1 isolation was performed by covering the rest of the window with a black silicone elastomer (Dow Corning Sylgard 170, Ellsworth adhesives). The light intensity level used was 0.45mw/mm^2^.

### Data analysis – widefield imaging

As in Wekselblatt et al. (2016), to analyze widefield fluorescence images while correcting for hemodynamic noise, 3:1 alternating blue and green frames were separately interpolated to produce a continuous image series at 10Hz. The fractional fluorescence change relative to the mean over the recording period (dF/F) was calculated for each pixel in each channel. The green channel was subtracted from the blue channel to correct for hemodynamic signals, giving a corrected fractional fluorescence intensity dF/F.

For periodic stimuli (Figure 3), we computed the amplitude and phase of the Fourier component of the dF/F signal at the stimulus frequency (0.1Hz). For each pixel, the Fourier phase corresponds to the retinotopic position that elicited the greatest visual response from the neural population at that cortical location. Retinotopic maps generated from widefield imaging were then used to target two-photon recordings to the cortical region within the cranial window representing the behavioral stimulus locations, either top or bottom, which were identified based on vasculature.

### Data analysis – two-photon imaging

Two-photon image data was first spatially aligned within the acquisition software, using phase correlation to estimate x-y translation. Cell body ROIs were extracted semi-automatically using published constrained non-negative matrix factorization algorithm, including neuropil correction and temporal deconvolution (Pnevmatikakis et al., 2016). Fractional fluorescence change (dF/F) was calculated for each cell, for both behavior and passively viewed stimuli.

Periodic mapping was performed immediately after two-photon behavioral sessions to confirm accurate targeting of the retinotopic location in V1 representing the stimulus location. To ensure we had estimates of the spatial receptive field centers for all cells, we calculated a smooth map of retinotopy based on the neuropil response. Next, we calculated the retinotopic positions of each individual neuron based on its position on the neuropil map. Two-photon retinotopic mapping data was then used to restrict analysis only to cells whose location in the retinotopic map overlapped with the locations of behavioral stimuli. Although the stimuli were 33° in diameter, only cells whose receptive field centers fell within a 24° diameter (‘centered’) were used for analysis.

Hierarchical clustering was performed on the response time-courses of imaged ‘centered’ cells for rewarded trials using Wards criterion (Matlab). For each training group, analyses of modulation due to locomotion and orientation selectivity were performed on all active cells for each cluster. For analysis of group differences of each cell class across training conditions, cell responses were averaged by individual sessions, and then session averages were combined among like training groups.

### Experimental Design and Statistical Analysis

Adult mice 2–6 months old, both male and female, were used in this study. Mice used for imaging experiments were a cross of GCaMP6s under the control of CRE (Jackson Labs 024742 (Wekselblatt et al., 2016)), and CaMK2-tTA (Jackson Labs 007004 (Mayford et al., 1996)). For optogenetic silencing of visual cortex, we crossed mice expressing Cre-dependent ChR2 (Jackson Labs 012569) and Pv-Cre (Jackson Labs 008069). A total of 15 animals were used for imaging experiments, with 5 mice in each training group (51 total sessions, 10,605 total cells). 7 animals were used for optogenetic silencing experiments (26 total sessions).

Statistical tests on cell responses were performed using the Kruskal-Wallis test, a nonparametric version of a one-way ANOVA. Multiple comparisons were made using Tukey’s honestly significant difference procedure. Significance testing for the fraction of active cells was performed in Matlab using a one-way ANOVA corrected for multiple comparisons. Unless otherwise noted, summary statistics are presented as means, with error bars representing standard error of the mean. Results were considered significant at p<0.05.

## Results

### A head-fixed paradigm to investigate sensorimotor learning in mice

For this study we used two-photon imaging of head-fixed mice to determine the effects of sensorimotor learning on V1 processing (Figure 1A). We compared cortical responses to visual stimuli in mice that were trained on a visual discrimination task (“trained”) with those of untrained mice (“naïve”). In addition, to test whether any changes observed were due to training or simply exposure, we imaged mice that had viewed the same stimuli over a similar number of sessions as the trained mice but were not trained on the discrimination rule (“experienced”).

To separate the acquisition of task structure (initiating trials and response) from learning of the visual discrimination itself, we first trained all experimental groups on a simple shaping task, based on luminance detection in which the mice had to move the Styrofoam ball left when the medial portion of the screen was light and right when the temporal portion of the screen was light. Once mice consistently performed >200 trials per session at over 90% performance (~1-2 weeks), they were assigned to one of the 3 experimental groups described above. Initial shaping allowed us to select for mice that were adept at self-initiating trials and responding in a 2-alternative forced choice (2-AFC) visual discrimination task, so that differences were due to learning the visual task and not to learning the trial structure.

We then trained mice to perform a spatial localization task, reporting whether a single square-wave grating patch appeared on the top or bottom of the monitor. Mice were trained to respond identically to grating patches of either horizontal or vertical orientation (Figure 1B). Animals assigned to the ‘trained’ group were rewarded for responding correctly based on the location, not the orientation, of the grating. Animals became proficient at the spatial discrimination task in 2-4 weeks. We performed imaging of the ‘trained’ group only after animals performed >200 trials per session at over 80% correct for 2-3 consecutive days. Animals in the ‘naïve’ group were imaged during their initial exposures to the grating stimuli (<8 sessions), immediately following mastery of the shaping task (Figure 1C, D). Due to this shaping, ‘naïve’ mice would self-initiate trials and respond at approximately the same frequency and time-course as the other two training groups (Figure 2A-C). Animals assigned to the ‘experienced’ group were presented with the same grating stimuli, but the rewarded response direction was random with respect to the stimulus. Thus, we exposed ‘experienced’ animals to the same visual stimuli as the ‘trained’ group, but without allowing learning of a specific discrimination rule. Animals in the ‘experienced’ group were given approximately the same amount of exposure to the grating stimuli as the ‘trained’ group before imaging occurred (1-2 months, >50 sessions, >10,000 trials).

**Figure 2.**
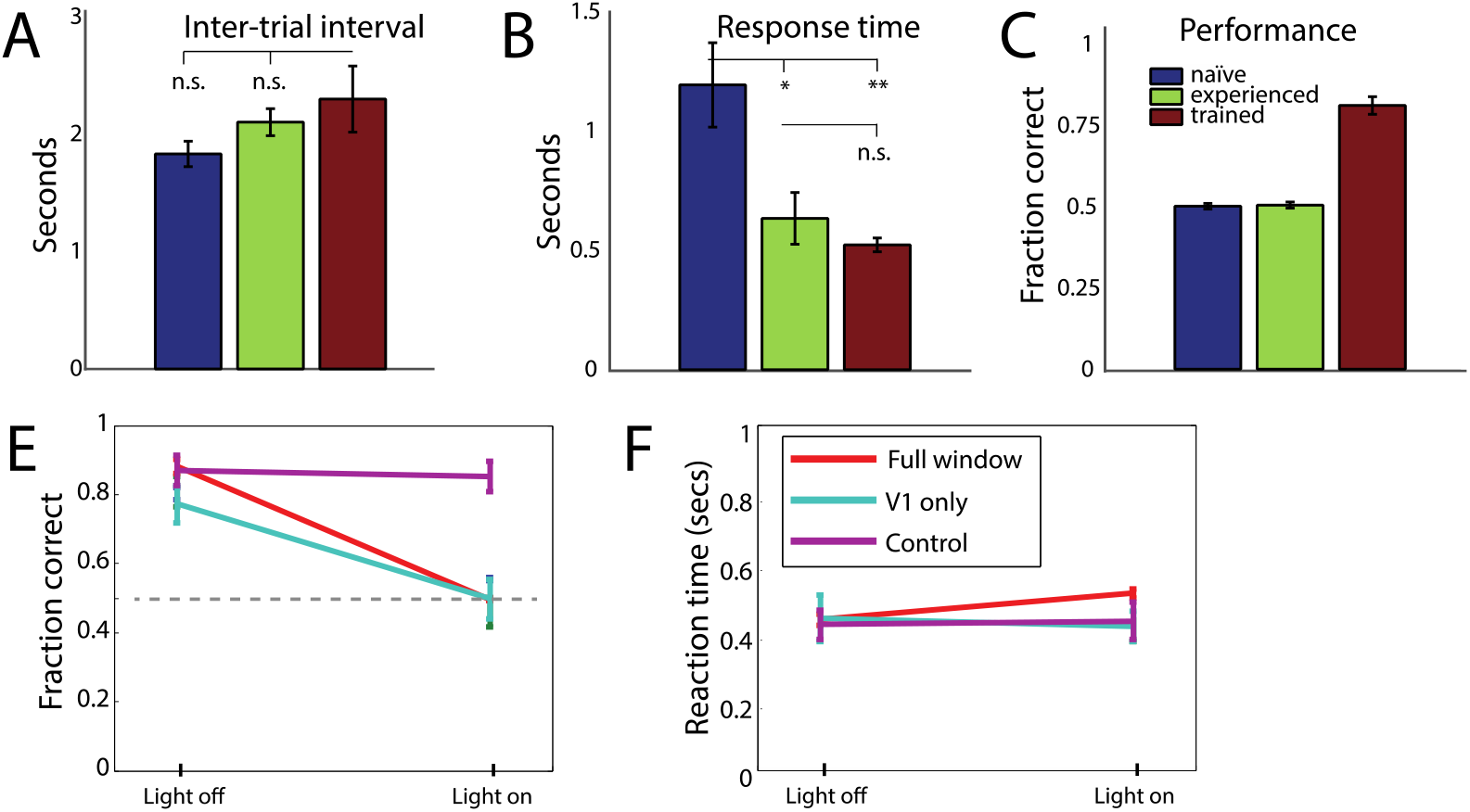
Task performance and dependence on visual cortex. **A, B)** Mean trial initiation times (A) and response times (**B)** for each training category, showing that all learned the task structure equally well. **C)** Mean performance in visual discrimination behavior for each group. **E**-**F)** Optogenetic silencing of visual cortex. **E)** Performance on spatial discrimination task is dependent on visual cortex. **F)** Mean reaction time with and without shutdown.

### Optogenetic silencing of visual cortex disrupts performance in the spatial location task

To determine whether visual cortex was necessary for the performance of the spatial discrimination task, we performed optogenetic inactivation of visual cortex in a separate group of animals. For this manipulation, we trained transgenic animals expressing Cre-dependent channelrhodopsin-2 (ChR2) in parvalbumin positive (Pv+) interneurons. Prior studies have shown that application of blue light in Pv+/ChR2 animals causes almost complete silencing of the excitatory population, by strongly activating inhibitory drive to excitatory cells through local GABAergic fast-spiking interneurons (Lien and Scanziani, 2013; Burgess et al., 2017)

After reaching criterion performance in the spatial location task, animals received blue light stimulation through a chronic cranial window on 20% of trials. Silencing was done either by illuminating the entire 5 mm cranial window (which includes several extrastriate visual areas) or restricting illumination to V1 by blocking access to the rest of the window using a black silicone elastomer (Poort et al., 2015). Light was delivered with a blue LED (470 nm, 0.45 mW/mm^2^) from stimulus onset until a behavioral response was registered. We found that performance in the spatial discrimination task was significantly worse on silencing trials compared to normal trials within the same session for animals expressing ChR2, in both the entire window (p=0.004, N=6 sessions) and V1 shutdowns (p=0.002, N=7 sessions). Performance was not affected for control animals that express only GCaMP6 (Figure 2D). Importantly, reaction time was unchanged for all groups, suggesting that this impairment in performance was not due to impairment of motor output (Figure 2E).

### Widefield mapping of visual cortex and targeting of stimulus location

To image the response characteristics of different excitatory neurons in V1, we used transgenic mice expressing a genetically encoded calcium indicator, GCaMP6s (Chen et al., 2013), under control of a CaMK2 driver line (Mayford et al., 1996). In these mice the majority of excitatory neurons in cortex express GCaMP6s (Wekselblatt et al., 2016). In order to target imaging of cortical calcium signals to the specific cortical locations within V1 that represent the behavioral stimulus positions, we performed widefield retinotopic mapping prior to our two-photon recordings (Figure 3). By mapping both azimuth and elevation using a moving window of periodic topographic noise as previously described in Wekselblatt et al. (2016), we generated maps of the preferred locations in visual space for the population activity of all the pixels within the 5 mm cranial window (Figure 3A-C). This allowed us to determine the specific cortical locations within V1 that represent the behavioral stimulus positions for each individual mouse using an overlay of vasculature landmarks (Figure 3A, D).

**Figure 3.**
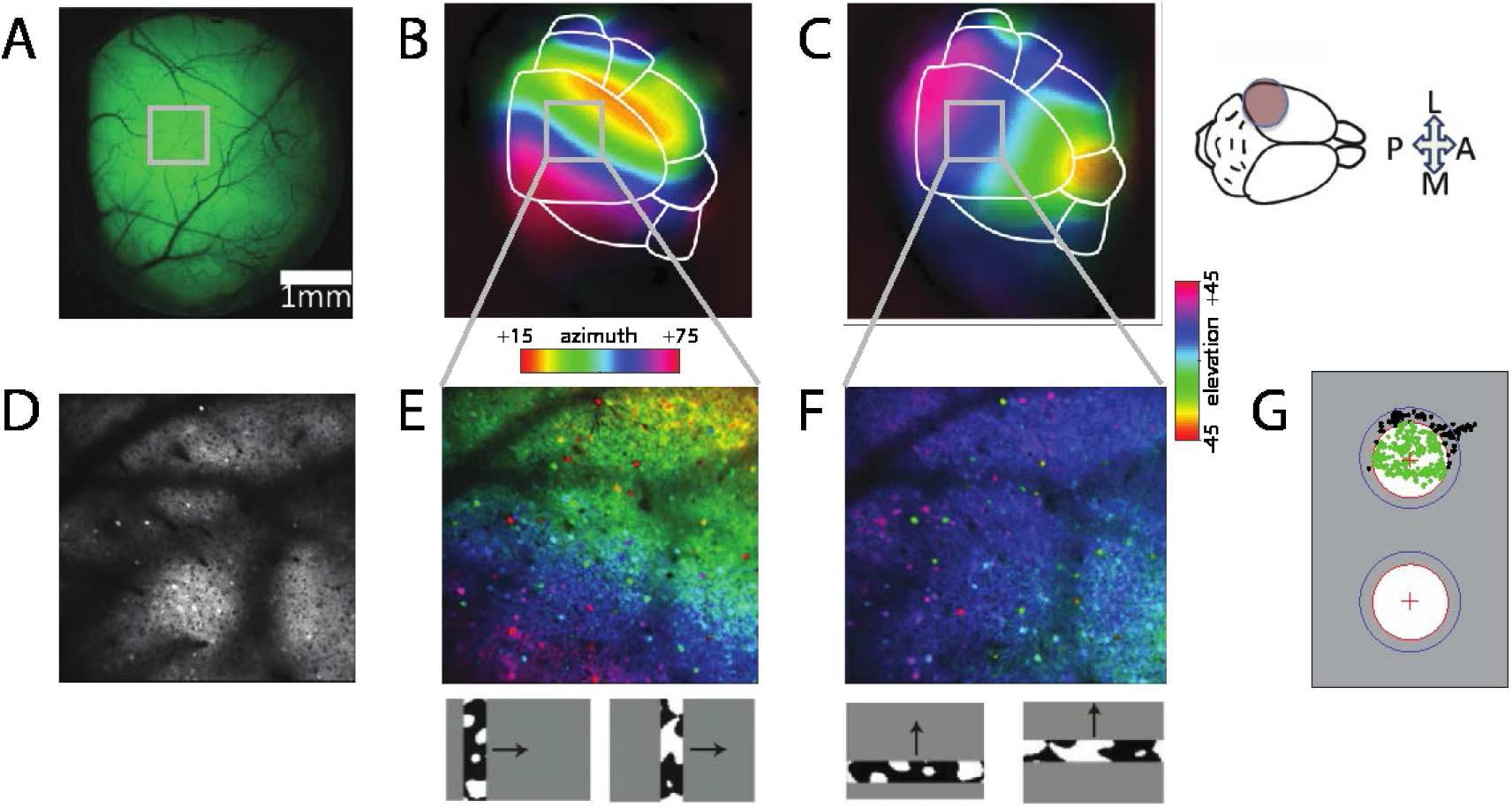
Targeted imaging of stimulus location in primary visual cortex (V1) **A)** Single photon illumination of GCaMP6-s expression in visual cortex through chronic cranial window. Inset shows location of 2-photon recoding shown in (D-F). Scale bar = 1mm, applies to (A-C). **B**-**C)** Widefield retinotopic mapping of azimuth **(B)** and elevation **(C)**, to functionally target 2-photon recordings to stimulus location. Colors represent position on monitor that elicited the greatest activity from each cortical location. **D)** 2-photon field of view in layer 2/3 of V1 (800um x 800um). **E**-**F)** 2-photon retinotopic mapping allows precise targeting of stimulus location in azimuth (E) and elevation (F). **G)** Receptive field centers for cells recorded from a single session. Blue circles denote stimulus locations (33 degree diameter). Inner red circles represent selection criteria (24 degree diameter) for ‘centered’ cells included in analysis. Green dots are included cells. Black dots are excluded cells. Inset at top right shows location of cranial window and orientation of animal for panels A-F. Inset at bottom shows schematic of retinotopic mapping stimulus for azimuth (B, E) and elevation (C, F).

To further confirm that the field of view (~800um x 800um) in our two-photon recordings was centered at one of the grating patch stimulus locations presented during behavior (top or bottom), mapping of the retinotopic position of each cell’s receptive field was performed following each behavior imaging session (Figure 3E, F). This additional mapping step allowed us to select only cells whose receptive field locations were well within the bounds of our behavioral grating stimulus for further analyses (Figure 3G). Although the grating stimulus was ~33° in diameter, only cells with receptive fields that fell within the central 24° diameter were used in subsequent analyses; this conservative bound was set so that only cells that were well-centered on the stimulus were included. One stimulus location was imaged per session, and the location imaged was alternated between the retinotopic location in V1 representing the top and bottom stimulus positions on each subsequent session. We then combined the responses across sessions to the stimulus position that corresponded to the receptive field location of the recorded cells for that session, which we refer to as the “retinotopically matched” location.

### Heterogeneity in layer 2/3 excitatory population

We next measured task-evoked activity in functionally defined regions of primary visual cortex while mice performed the behavioral tasks described above. Recordings were made at various depths ranging from 100 to 200 um below the surface of the brain, in layer 2/3 of V1, during both behavior and passive viewing. We could simultaneously observe the activity of hundreds of imaged excitatory neurons. Figure 4A shows the time course of activity of all (N=10605) recorded neurons with receptive fields within the retinotopic locations of the stimuli, during correct responses for each of the stimulus conditions.

**Figure 4.**
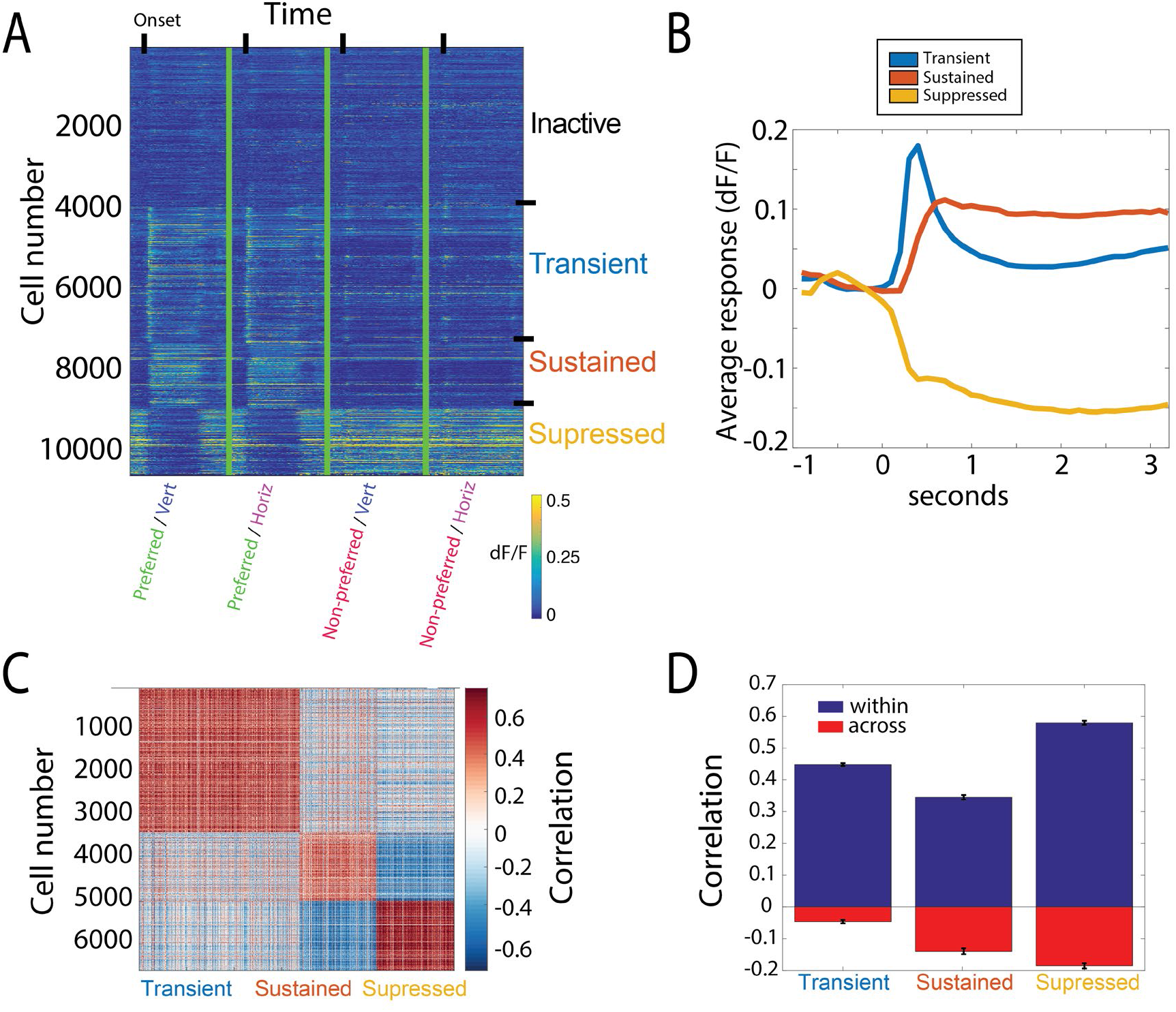
Hierarchical clustering of response time-course reveals striking heterogeneity among excitatory neurons. **A)** Average response (dF/F) of imaged neurons in response to each of the four stimuli used for behavior (correct trials only). Responses sorted according to cluster identity based on time-course (inactive, transient, sustained, suppressed). **B)** Average evoked response time-course for each functional cell class, determined by cluster identity. Average from correct trials at the retinotopically matched location only, pooled across orientations. T=0 represents stimulus onset. **C)** Correlation of the mean timecourse of response between all pairs of recorded neurons, following clustering, demonstrating high correlation within blocks of clustered cells, with weaker correlations across cells in different clusters. **D)** Mean correlation for all pairs of cells within a cluster (blue), and pairs across a given cluster and the other two clusters (red).

Hierarchical clustering based on the response time-courses of these cells revealed striking heterogeneity across the population. Three distinct response types were found among neurons activated during behavior, which we term “transient”, “sustained,” and “suppressed” (Figure 4B). Transient cells exhibited a large and brief response at the onset of the visual stimulus that quickly decayed. Sustained responding cells showed a slower rise to peak and maintained activity until the offset of the visual stimulus. Suppressed cells had high baseline activity and were suppressed during the time that the visual stimulus was present.

These classes each showed a high correlation in the timecourse of their response within a class, with much weaker or negative correlations across classes (Figure 4C, D). This indicates that the clustering defines three groups with distinct timecourses in aggregate. As shown below, these groups also differ in other aspects of their response. We used these classifications to group cells for subsequent analyses. In addition, cells that had less than a 0.02 standard deviation in the dF/F of their response during behavior were deemed ‘inactive,’ and were excluded from clustering and subsequent analyses. Changing this criterion by a factor of two in either direction did not significantly affect results.

### Different functional cell classes show distinct selectivity and state dependence

To allow us to assess the effects of task-engagement on V1 processing, and to measure orientation selectivity and preference of recorded neurons, we introduced an additional passive viewing stimulus. Following the behavior session, the mice were presented with static grating patches of four evenly spaced orientations (0, 45, 90, 135 deg), adding two orientations not shown in previous stimuli. The water-spout was removed following the behavior session, and passive stimuli were presented in a pseudo-randomized order for 1 second each, with a 1.5 second inter-stimulus-interval.

The distribution of orientation selectivity differed significantly (p<0.001) among these cell classes (Figure 5A-D). Sustained and transient cells had relatively high orientation selectivity (mean OSI = 0.55 +/− 0.01, and 0.48 +/− 0.01 respectively). In contrast, suppressed cells showed very little orientation preference (mean OSI = 0.28 +/− 0.01), as has been reported for suppressed-by-contrast cells in previous studies (Tailby et al., 2007; Niell and Stryker, 2010).

**Figure 5.**
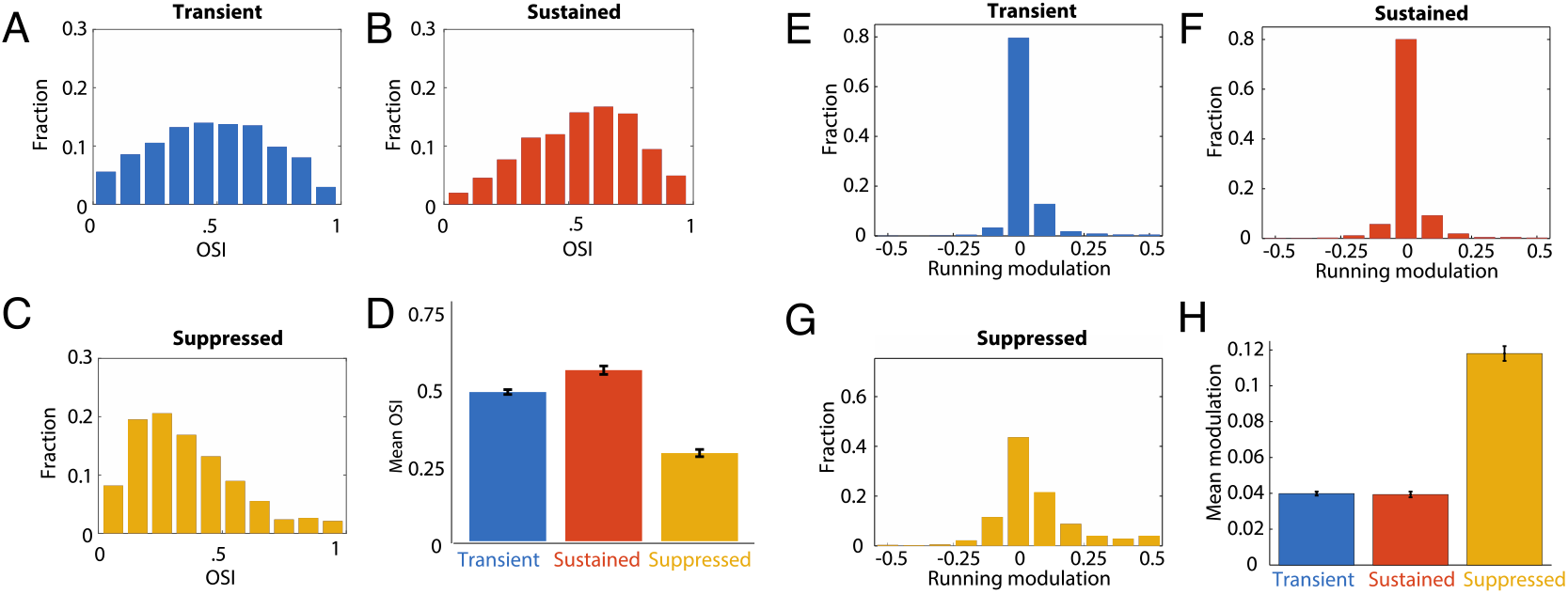
Distinct functional cell classes exhibit unique differences in orientation selectivity and state modulation. **A**-**C)** Distribution of orientation selectivity (circular variance) for transient cells (A), sustained cells (B), and suppressed cells (C). **D)** Average orientation selectivity for cells of each functional class. **E**-**G)**. Distribution of modulation by running of spontaneous activity (S) for transient cells (E), sustained cells (F), and suppressed cells (G). **H)** Average running modulation for each functional cell class, showing suppressed cells show greatest modulation of spontaneous activity by running. Modulation of spontaneous activity by locomotor state de_ned by (Srun - Srest) / (Srun + Srest)

Furthermore, these cell classes were differentially modulated by locomotion (Figure 5E-H). Locomotion alters the gain of visual responses in many upper layer cells in mouse V1 (Niell and Stryker, 2010; Vinck et al., 2015; Reimer et al., 2014). We tracked locomotion during passive viewing sessions and examined its effect on the response profiles of the different cell classes. Suppressed cells showed a much greater modulation of baseline firing rate with locomotion than either the transient (p<0.001) or sustained (p<0.001) cell classes (Figure 5E-H), similar to previously reported suppressed-by-contrast cells (Niell and Stryker, 2010). The difference in response dynamics during behavior, passive receptive field properties, and level of modulation by behavioral state demonstrates that there is significant diversity in distinct sub-populations of layer 2/3 excitatory neurons in V1.

### Training affects proportion of responsive cells

To investigate whether there were long term changes in response properties of layer 2/3 cells as a result of learning or repeated exposure, we compared the fraction of responsive neurons, defined by having a standard deviation in dF/F greater than 0.02 during behavior, between the behavioral training groups (trained, naïve, and experienced). The proportion of responsive cells was smaller in ‘trained’ (26.6 +/− 3.0%) and ‘experienced’ (30.5 +/− 2.7%) animals relative to the ‘naïve’ group (37.2 +/− 2.6%) (p=0.044 and p=0.001 respectively). The reduction in responsive cells was due almost entirely to the decreased proportion of transient responses in both the ‘trained’ and ‘experienced’ groups compared to the naïve group (Figure 6), although this was only significant for the trained group (p=0.03). There were no significant changes in the number of sustained or suppressed cells between groups (p=0.12 and p=0.44 respectively).

**Figure 6.**
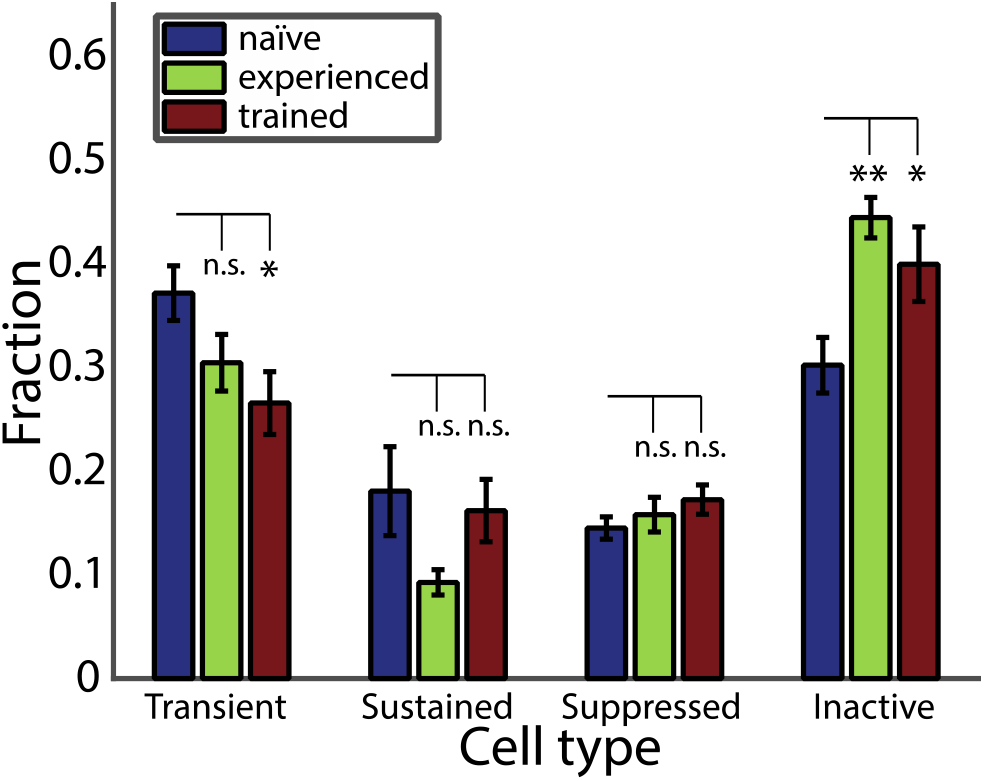
Task training and exposure result in a reduction of the fraction of active cells in layer 2/3 of V1. Fraction of active cells during behavior for each functional cell class and each training condition. Average is calculated across fractions in each individual session.

Next, we examined which cell types contributed to the diminished responses in learned and experienced animals. Both the ‘trained’ and ‘experienced’ groups showed a decreased proportion of transient responding cells compared to the ‘naïve’ group (Figure 6), although this was only significant for the trained group (p=0.03). We did not observe a significant change in the number of sustained or suppressed cells between groups (p=0.12 and p=0.44 respectively). This dissociation suggests a cell-type specific effect of learning versus extended exposure.

Despite the changes observed in the proportions of active cells, we did not find evidence that behavioral training affected response magnitude or time-course of activity within the responsive cells, for any of the cell classes across training groups (Figure 7). Of the cells that were active during the behavior, temporal dynamics (Figure 7A-C) and the peak amplitude (Figure 7D) of the response were nearly identical across training conditions for all three cell classes. This finding is consistent with previous studies which show learning related changes primarily affect the fraction of responsive cells, but not the magnitude of the responses in active neurons (Makino and Komiyama, 2015; Chu et al., 2016). While we did not perform longitudinal imaging to follow individual neurons over time, these findings suggest that a significant portion of the transient cells become unresponsive as a result of repeated exposure.

**Figure 7.**
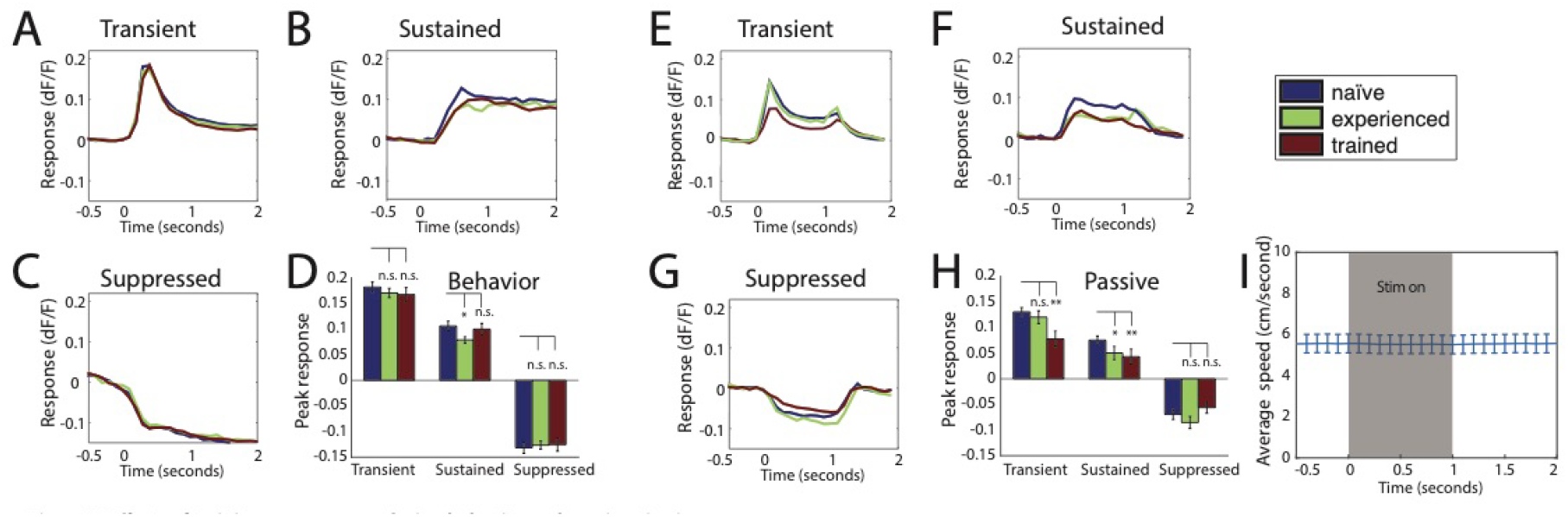
Effects of training on responses during behavior and passive viewing. **A**-**C)** Average timecourse of evoked activity during behavior for each cell class, by training condition. D) Average response at peak timepoint for each cell class and training group. Transient peak = 200-300ms, sustained/suppressed peak = 800-900ms. **E**-**G)** Average timecourse of evoked activity during passive viewing for each cell class, by training condition. **H)** Average response at peak timepoint for each cell class and training group. **I)** Average running speed during stimulus presentation period for passive viewing.

### Training condition affects active/passive modulation

We next examined whether training condition (‘trained’, ‘naïve’, or ‘experienced) had an effect on visually evoked responses to the grating stimuli when presented passively. To achieve this, we compared the activity evoked during the passive viewing sessions described above for measuring orientation selectivity. To control for the differences in the observed proportion of active cells among training groups, we restricted this analysis to cells that were active during behavior. Passive responses from animals in the ‘experienced’ group were not different from those in the ‘naïve’ group. In contrast, the calcium signals of both transient (p=0.005) and sustained (p=0.001) cells for animals in the ‘trained’ group were smaller than those in naïve animals (Figure 7E-H). Importantly, we did not observe a change in temporal dynamics between groups during passive viewing, verifying that the classification into groups based on temporal dynamics was robust in all conditions of these experiments. Furthermore, there was no locomotor activity locked to stimulus onset during passive viewing (Fig 7I), suggesting that the characteristic timecourse of each group is not due to locomotion. Note that the time-course differ between passive viewing and the behavioral task due to stimulus presentation – in passive viewing stimuli were presented for 1 sec, whereas during behavior they remained on the screen until 1.5 sec after the animal’s response, resulting in longer stimulus presentation.

To determine how learning affected the level of modulation by task engagement, we compared active and passive responses to the grating stimuli for each cell class within the three training groups. We found that responses during active behavior were significantly larger than the responses to the passive viewing of the same stimuli for all cell classes within all training groups (range of p-values = 7.7×10^−5^ to 0.0016, across nine groups) (Figure 8A-C). This difference between active and passive responses was especially pronounced for the group of animals who had been trained on the spatial discrimination (Figure 8B). To quantify this observation, we calculated a task modulation index ((task − passive) / (task + passive)) for each cell. Figure 8D shows that both transient and sustained cell classes in animals trained in the spatial location task had significantly higher task modulation indices compared to ‘naïve’ animals (transient: p = 0.014, sustained: p = 0.017, suppressed: p = 0.160). Task modulation in ‘experienced’ animals, on the other hand, was similar to that of ‘naïve’ animals. Thus, in addition to cell-type specific changes in the proportions of active cells with learning, the modulation of responses due to task-engagement depended on the specific training history.

**Figure 8.**
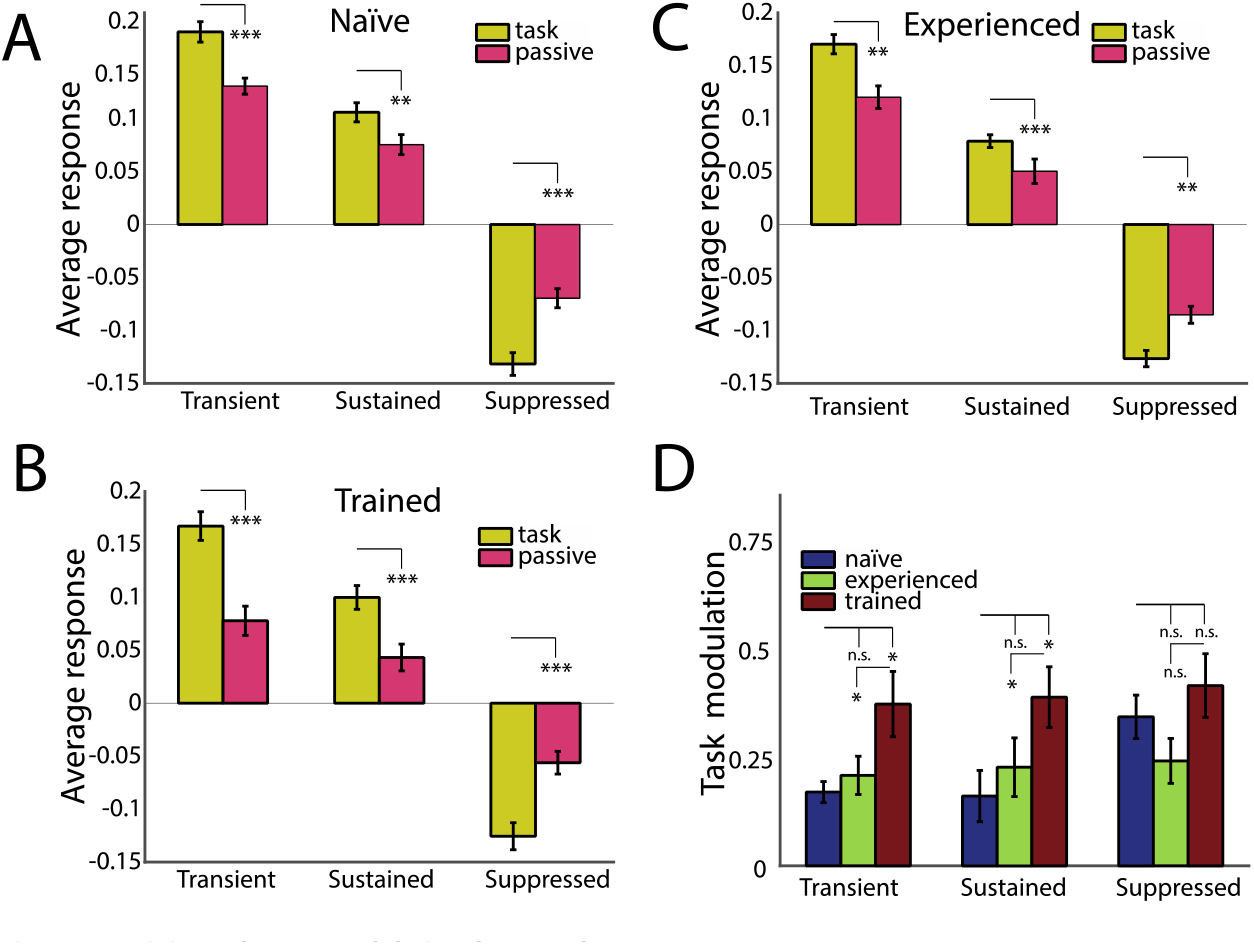
Training enhances modulation due to task engagement. **A**-**C)** Task evoked responses are enhanced relative to passive responses to the same stimuli for all training groups and cell classes. Responses evoked to grating stimuli during behavior and passive viewing for naïve group (A), trained group (B), and experienced group (C). **D)** Task modulation is enhanced by learning. Trained animals show greater modulation for transient and sustained cell classes. Task Modulation Index = (Rtask − Rpass) / (Rtask +Rpass).

## Discussion

In this study we determined how different excitatory neuron types are modulated by learning and task engagement. We show that excitatory cells in layer 2/3 can be characterized into three functionally distinct sub-populations based on time-course of response to a visual stimulus: ‘transient’, ‘sustained’ and ‘suppressed’. Furthermore, we show that these cell classes exhibit different selectivity and modulation by behavioral state. Experience and learning reduce the fraction of cells responsive to visual stimuli used in the behavior task. Notably, we find a specific reduction in the transient population in trained animals. Additionally, we find that trained animals show greater modulation due to active task-engagement compared with naïve and experienced animals for both the transient and sustained cell classes.

Previous research has shown similar decreases in population activity after learning in both human and animal studies (Mruczek and Sheinberg, 2007; Anderson et al., 2008; Woloszyn and Sheinberg, 2013). Additionally, a reduction in active neurons over training has been observed in several brain areas including auditory cortex, olfactory bulb, motor cortex and visual cortex suggesting a potentially shared mechanism across sensory systems (Otazu et al., 2009; Peters et al., 2014; Kato et al., 2015; Makino and Komiyama, 2015; Chu et al., 2016). This shift in the fraction of active neurons may be an effect of repeated exposure to the same stimuli over the course of thousands of trials, or may reflect a more general design principle of sensorimotor learning in which sensory cortex acquires efficient coding after training (Makino et al., 2016). Primary visual cortex has many downstream targets including extrastriate cortex, posterior parietal cortex, retrosplenial cortex, striatum and amygdala. The reduction in the fraction of behaviorally responsive neurons over training may represent an early mechanism for gating information flow to the appropriate target areas needed to execute the proper behavioral output. This is an intriguing idea given that cortico-cortical neurons in V1 have been shown to convey specific information matched to the preferences of the downstream recipient area (Zeki and Shipp, 1988; Nassi and Callaway, 2009; Glickfeld et al., 2013; Glickfeld and Olsen, 2017).

A number of recent studies have shown that motor output and other state variables are reflected in neural activity, even in primary visual cortex (Niell and Stryker, 2010; Keller et al., 2012; Musall et al., 2018; Stringer et al., 2018). It is therefore possible that some aspects of the response timecourse, and changes with learning, could reflect the motor output associated with task performance. Indeed, the high baseline activity of the suppressed group is strongly modulated by locomotion. However, we do not expect that this explains our classification or differences in learning across groups, as the same response types were observed even in passive viewing, where there was no stimulus-evoked motor output (Figure 7). It is possible, though, that further distinctions among the excitatory population, and their changes with learning, may arise if cells are classified by the motor signals they encode.

Our study provides the first report of sub-divisions within the cortical excitatory cell population that are differentially affected by learning and context. This characterization is an important step in determining the specific mechanisms by which information flow is routed to the correct brain areas for action. Local inhibitory neurons are also likely involved in this learning-related reduction of excitatory responses. In primary visual cortex, somatostatin (SOM) interneurons have been implicated in gating top-down effects of task engagement (Makino and Komiyama, 2015), and in auditory cortex both SOM+ and Pv+ interneurons have been shown to play a role in perceptual learning by providing context dependent synaptic inhibition (Kato et al., 2015; Kuchibhotla et al., 2016). While context-dependent changes in inhibitory activity primarily affect local processing, changes in the excitatory cell population, especially in layer 2/3, will have effects on both local and long-range synaptic partners and could thus function to gate the information sent to downstream areas (Zeki and Shipp, 1988; Nassi and Callaway, 2009; Glickfeld et al., 2013; Glickfeld and Olsen, 2017). It will be important for future studies to determine how these excitatory cell classes are connected with and affected by different interneuron sub-types and long-range projections to and from other brain areas.

It is important to note that although the ‘experienced’ group was not trained on an explicit discrimination, this does not imply that no learning was involved in their performance. Due to the structure of the task, it is likely that these animals adopt a detection strategy, rather than a discrimination strategy, learning to move the ball and lick for reward when the stimulus appears even though there is no predictive power in the content of the stimulus. This may contribute to the similarities we see between ‘experienced’ and ‘trained’ animals, in terms of the reduction of responsive neurons during behavior. Replication of our findings with similar and different task demands should be done to address whether the observed changes are task-specific or a general mechanism of learning.

We also note that because we did not perform chronic imaging of the same cells, we cannot say what changes occurred to individual neurons over learning. Although we report population differences rather than changes in the same cells over time, our results are consistent with previous longitudinal studies on changes in the responses of single neurons during learning (Kato et al., 2015; Makino and Komiyama, 2015; Chu et al., 2016). Future longitudinal studies should investigate the time-course of the reduction of responses in individual neurons, and how this time-course compares with that of increased behavioral proficiency.

The excitatory population in cortex labeled by the CaMK2 promoter includes roughly 80% of the neurons in cortex (Mayford et al., 1996; Wekselblatt et al., 2016). Here we show that this population is diverse and consists of at least three cell types, as defined by their functional properties (response timecourse). The functional differences observed here might reflect different inputs to the identified cell classes, potentially originating from different retinal ganglion cell types (Solomon et al., 2002; Tailby et al., 2007). On the output side, these populations may represent parallel processing streams analogous to the dorsal and ventral streams identified in humans and primates, which have distinct projection patterns to downstream areas and are specialized for processing different information (Ungerleider and Mishkin, 1982; Nassi and Callaway, 2009; Wang et al., 2012). Future studies should be able to identify these cell types using molecular or genetic markers, projection patterns, or anatomical reconstruction to help determine their role in cortical circuits underlying learning and behavior.

## Acknowledgments

This work was supported by NIH T32HD007348 (J.B.W.), NIH F31EY025459 (J.B.W.), NIH R01EY023337 (C.M.N.), and ONR N00014-16-1-3154 (C.M.N.). We thank Dr. Matt Smear, Dr. Michael Stryker, Dr. Doris Tsao, and members of the Niell lab for helpful discussions and comments on the manuscript.

## Author contributions

J.B.W. and C.M.N. conceived and designed the study. J.B.W. performed experiments. J.B.W. and C.M.N. analyzed data and prepared the manuscript.

## Declaration of Interests

The authors declare no competing interests.

